# Glucocorticoids rapidly modulate Ca_V_1.2-mediated calcium signals through Kv2.1 channel clusters in hippocampal neurons

**DOI:** 10.1101/2023.12.24.573220

**Authors:** Di Wan, Tongchuang Lu, Chenyang Li, Changlong Hu

**Author notes:** These authors contributed equally to the work (Di Wan, Tongchuang Lu). **Corresponding author:** Changlong Hu.

## Abstract

The precise regulation of Ca^2+^ signals plays a crucial role in the physiological functions of neurons. Here, we investigated the rapid effect of glucocorticoids on Ca^2+^ signals in hippocampal neurons. In cultured hippocampal neurons, glucocorticoids inhibited the spontaneous somatic Ca^2+^ spikes generated by Kv2.1-organized Ca^2+^ microdomains. Furthermore, glucocorticoids rapidly reduced the cell surface expressions of Kv2.1 and Ca_V_1.2 channels in hippocampal neurons. In HEK293 cells transfected with Kv2.1 alone, glucocorticoids significantly reduced the surface expression of Kv2.1 with little effect on K^+^ currents. Glucocorticoids inhibited Ca_V_1.2 currents but had no effect on the cell surface expression of Ca_V_1.2 in HEK293 cells transfected with Ca_V_1.2 alone. Notably, in the presence of wild-type Kv2.1, glucocorticoids caused a decrease in the surface expression of Ca_V_1.2 channels in HEK293 cells. However, this effect was not observed in the presence of non-clustering Kv2.1S586A mutant channels. Live cell imaging showed that glucocorticoids rapidly decreased Kv2.1 clusters on the plasma membrane. Correspondingly, western blot results indicated a significant increase in the cytoplasmic level of Kv2.1, suggesting the endocytosis of Kv2.1 clusters. Glucocorticoids rapidly decreased the intracellular cAMP concentration and the phosphorylation level of PKA in hippocampal neurons. The PKA inhibitor H89 mimicked the effect of glucocorticoids on Kv2.1, while the PKA agonist forskolin abrogated the effect. In conclusion, glucocorticoids rapidly regulate Ca_V_1.2-mediated Ca^2+^ signals in hippocampal neurons by promoting the endocytosis of Kv2.1 channel clusters through reducing PKA activity.

**Significance Statement:** The rapid non-genomic effect of glucocorticoids on the central nervous system is not fully understood. Ca_V_1.2-mediated Ca^2+^ signaling microdomains control somatic Ca^2+^ signals and regulate excitation-transcription coupling in hippocampal neurons. Here, we demonstrate that glucocorticoids rapidly inhibit Ca_V_1.2-mediated somatic Ca^2+^ spikes in hippocampal neurons. Glucocorticoids reduce the surface expression of Kv2.1 clusters but do not affect the surface expression of non-clustering Kv2.1. Moreover, glucocorticoids induce the endocytosis of Ca_V_1.2 channels through wild-type Kv2.1. However, glucocorticoids cannot induce the endocytosis of Ca_V_1.2 channels through Kv2.1S586A mutant channels, which cannot form clusters. This study sheds light on the intricate interplay between glucocorticoids, Kv2.1 channels, and Ca_V_1.2 channels, unraveling a dual mechanism that influences both overall Ca^2+^ signaling and the intricate organization of neural microdomains.

## Introduction

Glucocorticoids, also known as stress hormones, are produced by the zona fasciculata cells of the adrenal cortex in a diurnal pattern. In the mammalian brain, glucocorticoids induce variable neuronal effects via both genomic and non-genomic signaling pathways. While the genomic pathway has been well studied, the rapid non-genomic effects of glucocorticoids, which usually occur within a few minutes, are still not fully understood. The hippocampus is the main effector of glucocorticoids in brain (1). The level of glucocorticoids shows a distinct circadian and ultradian rhythm in brain (2, 3). Recent studies have shown that the ultradian rhythmicity of glucocorticoids is critical in regulating normal emotional and cognitive responses in humans (4). Glucocorticoids may modulate neural function through rapid non-genomic effects during these fluctuations.

The Ca^2+^ entry through L-type calcium channels, mainly Ca_V_1.2 and Ca_V_1.3 channels, is essential for the normal functioning of various mammalian brain neurons(5, 6). Even through both mRNA expression of Ca_V_1.2 and Ca_V_1.3 has been suggested in the hippocampus, L-type Ca^2+^ current in hippocampal neurons may be predominantly mediated by Ca_V_1.2 channels(7–10). Accordingly, the hippocampal pyramidal neurons in Ca_V_1.2 knockout mice displayed a nearly complete loss of L-type Ca ^2+^ currents, and Ca_V_1.2 but not Ca_V_1.3 knockout mice showed deficits in hippocampal long-term potentiation and memory formation(11–13).

The family of Kv2 channels, consisting of Kv2.1 and Kv2.2, plays a vital role in controlling neural excitability(14, 15). The Kv2.1 channel is widely expressed throughout the mammalian brain, and is the major conductor of rectifying potassium currents in hippocampal neurons(16, 17). Kv2.1 channels in cell membrane exist in two forms: clustered and nonclustered(18). Nonclustered Kv2.1 channels conduct K^+^ normally and are critical for maintaining repetitive firing of neurons(19, 20). Clustered Kv2.1 channels are barely conductive and serve as a platform to organize neuronal endoplasmic reticulum/plasma membrane (ER/PM) junctions(21–25). ER/PM junctions are critical for lipid and Ca^2+^ homeostasis in eukaryotic cells(26, 27), and are particularly abundant in the soma of mammalian brain neurons(28). Recent studies have shown that clustered Kv2.1 channels recruit Ca_V_1.2 to somatic ER/PM junctions to form Ca^2+^ signaling microdomains. These microdomains control somatic Ca^2+^ signals and regulate excitation-transcription coupling in hippocampal neurons(29, 30).

In this study, we investigated that the rapid effects of glucocorticoids on Ca^2+^ signals in hippocampal neurons. Our results show that glucocorticoids reduce cell surface expression of Ca_V_1.2 by promoting endocytosis of clustered Kv2.1. This, in turn, inhibits somatic Ca^2+^ sparklets in hippocampal neurons through the PKA signaling pathway.

## Results

### Glucocorticoids reduce somatic spontaneous Ca^2+^ signals and Kv2.1 clusters in cultured hippocampal neurons

First, we investigated the effect of glucocorticoids on somatic spontaneous Ca^2+^ signals in cultured hippocampal neurons under internal reflection fluorescence (TIRF) microscopy. The fluorescent calcium indicator Cal-590 AM was used to detect the spontaneous somatic Ca^2+^ signals in cultured hippocampal neurons. Bath application of 1 μM corticosterone for 5 min significantly reduced the frequency and amplitude of spontaneous somatic Ca^2+^ sparks in hippocampal neurons, with no effect on the spark width (Fig. 1A-C; Supplemental Video S1).

**Figure 1.**
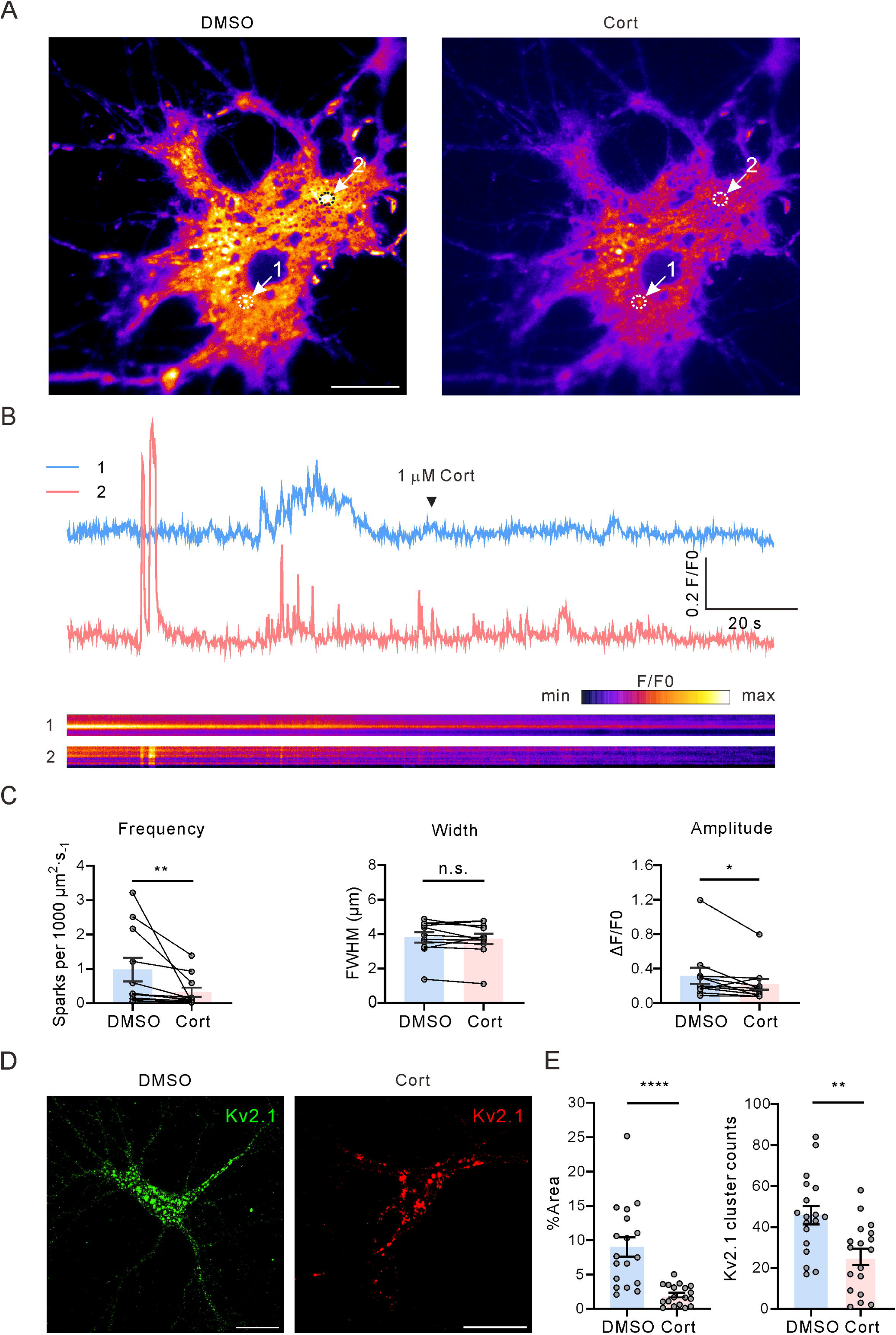
Effect of glucocorticoids on spontaneous Ca^2+^ sparks in cultured rat hippocampal neurons. (**A)** Representative TIRF images showing somatic Ca^2+^ sparks in hippocampal neurons (loaded with Cal-590 AM) under the vehicle control (0.01% DMSO) condition and subsequently in the presence of 1 μM corticosterone (Cort) in the same hippocampal neuron. Arrows indicate regions of interest (ROIs) where spontaneous Ca^2+^ signals were detected (scale bar: 10 μm) (Supplemental Video S1). **(B)** Fluorescence intensity traces (upper panels) and kymographs (lower panels) corresponding to the two ROIs indicated in panel A. **(C)** Statistics for the frequency, full width at half maximum (FWHM) and amplitude of all spatially distinct localized Ca^2+^ sparks recorded from hippocampal neurons before and after 1 μM corticosterone treatment using a two-tailed paired *t*-test (n = 11). **p = 0.0065; *p = 0.0461; n.s., not significant (p = 0.4463). Each point corresponds to a single neuron. **(D)** Representative examples of immunofluorescence images showing the Kv2.1 channel clusters in rat hippocampal neurons with treatment of 1 μM corticosterone or an equal volume of DMSO (scale bar: 20 μm). **(E)** Statistics for the percent area of Kv2.1 clusters and the number of Kv2.1 clusters in the somatic region of cultured hippocampal neurons from D using a two-tailed unpaired *t*-test (n = 18). ****p < 0.0001, **p = 0.0018.

Spontaneous somatic Ca^2+^ signals in hippocampal neurons are usually generated at Kv2.1 cluster-associated ER-PM junction sites, where Kv2.1 recruits Ca_V_1.2 to form local Ca^2+^-release microdomains(29). Therefore, we investigated whether corticosterone regulates Ca^2+^ signaling via Kv2.1 clusters. Kv2.1 clusters with diameter more than 0.5 μm were calculated (31). Treatment with 1μM corticosterone for 5 minutes reduced Kv2.1 clusters in the soma of hippocampal neurons (Fig. 1D). This suggests that corticosterone caused Kv2.1 declustering or endocytosis. To further investigate the mechanism by which glucocorticoids reduce somatic spontaneous Ca^2+^ signals in cultured hippocampal neurons, we tested the effect of corticosterone on Ca^2+^ and K^+^ channel currents in cultured hippocampal neurons.

### Glucocorticoids rapidly inhibit calcium and potassium channel currents in hippocampal neurons

Since the predominant calcium channel currents in hippocampal neurons are conducted by L-type Ca_V_1.2 channels (12), Ba^2+^ was used as the charge carrier to minimize Ca^2+^-induced inactivation. Whole-cell Ba^2+^ currents in cultured hippocampal neurons were elicited by a 100-ms depolarizing pulse to 0 mV from a holding potential of −90 mV at 10 s intervals. Bath application of 1μM corticosterone significantly inhibited the Ba^2+^ currents in hippocampal neurons (Fig. 2A). The inhibitory effect of corticosterone on *I*_Ba_ began quickly and reached a maximum effect around 5 min (Fig. 2B). Kv2.1 channels are the major contributor of delayed rectifier potassium currents (*I*_k_) in hippocampal neurons (16). *I*_k_ in hippocampal neurons was elicited by a 200-ms depolarizing pulse to +40 mV from a holding potential of −80 mV at 10 s intervals. 1μM corticosterone had a small inhibitory effect on *I*_k_ in hippocampal neurons (precent inhibition: 14.3 ± 2.9%, Fig. 2C). Previous studies have reported that Kv2.1 de-clustering is usually accompanied by a large hyperpolarizing shift in the steady-state activation of *I*_K_ in hippocampal neurons (32, 33). We found that extracellular application of corticosterone did not alter the steady-state activation of *I*_k_ in hippocampal neurons (Fig. 2D). This suggests that corticosterone may cause Kv2.1 cluster endocytosis instead of de-clustering. Therefore, we tested the effect of corticosterone on surface expression of Kv2.1 channels.

**Figure 2.**
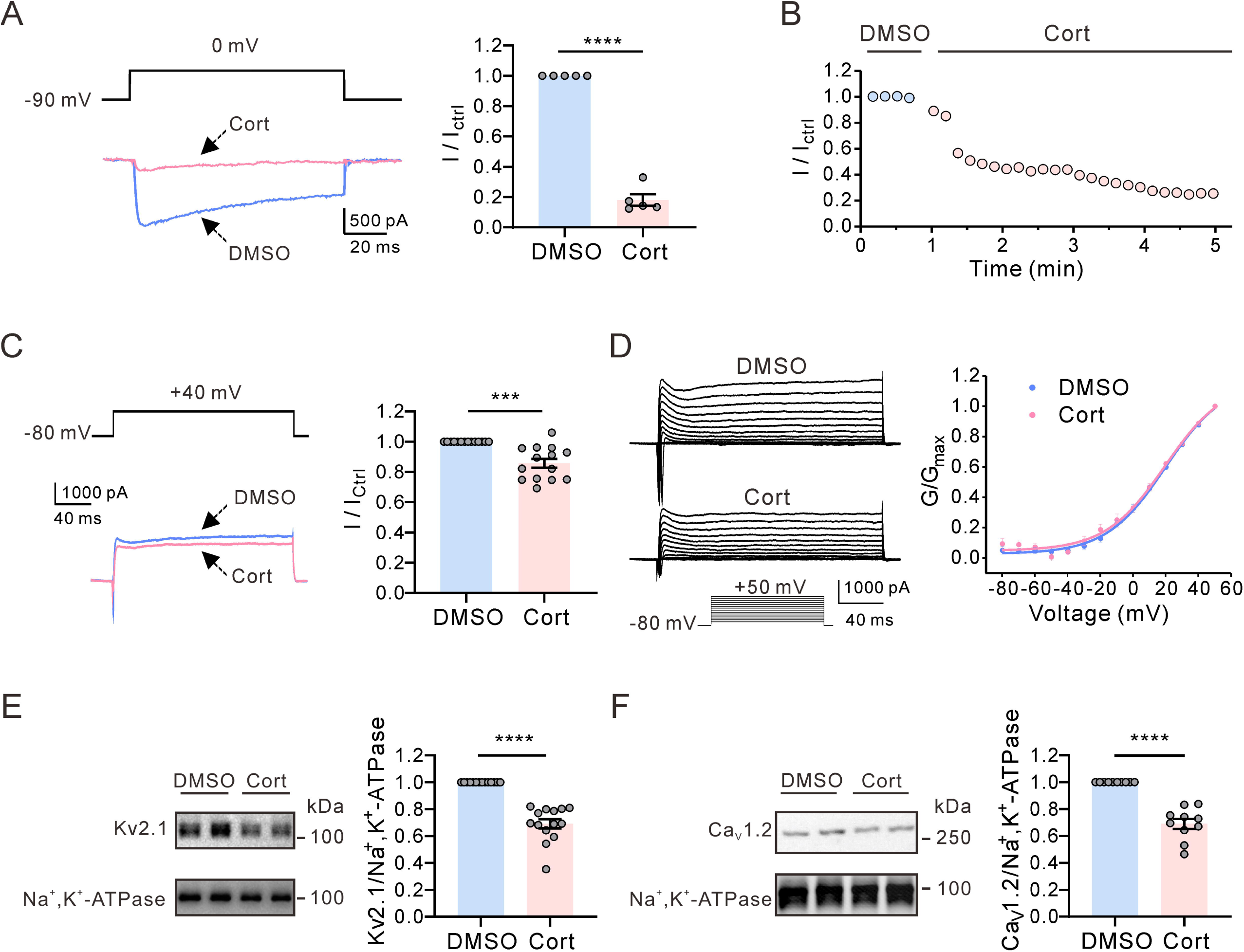
Effect of glucocorticoids on Ca^2+^ and K^+^ channel currents, and membrane protein expressions in cultured hippocampal neurons. **(A)** Left, representative *I*_Ba_ traces induced by a depolarization pulse from −90 to 0 mV under the vehicle control (0.01% DMSO) condition and subsequently in the presence of 1μM corticosterone in the same hippocampal neuron. Right, statistical analysis of the effect of corticosterone on *I*_Ba_ using a two-tailed paired *t*-test (n = 5). ****P = 2.8E-5. **(B)** The time course of the *I*_Ba_ inhibition by corticosterone. **(C)** Left, representative current traces show the inhibitory effect of 1μM corticosterone on *I*_K_. Right, statistics for the amplitude of *I*_K_ from Left using a two-tailed paired *t-*test (n = 14). ****P = 3.2E-4. **(D)** Left, representative *I*_K_ traces induced by a depolarization pulse from −80 to +50 mV under the vehicle control condition and subsequently in the presence of 1μM corticosterone in the same hippocampal neuron. Right, plot of *I*_K_ activation curves in the DMSO (n = 9 for each data point; blue) and corticosterone-treated (n = 9 for each data point; pink) groups. **(E)** Left, representative western blot of the Kv2.1 channel surface expression after 10-min corticosterone treatment (1μM) in the extracellular solution in cultured hippocampal neurons. Right, statistics from 14 independent experiments using a two-tailed unpaired *t*-test. **** P = 2.2E-9. Na^+^-K^+^ ATPase as a membrane protein loading control. **(F)** Left, representative western blot of the Ca_V_1.2 channel surface expression after 10-min corticosterone treatment (1μM) in the extracellular solution in cultured hippocampal neurons. Right, statistics from 10 independent experiments using a two-tailed unpaired *t-*test. ****P = 1.7E-7.

### Glucocorticoids reduce membrane protein expressions of Kv2.1 and Ca_V_1.2 in cultured hippocampal neurons

Treatment of cultured hippocampal neurons with 1 μM corticosterone for a duration of 10 minutes resulted in a significant reduction in the cell surface expression of Kv2.1 channels (Fig. 2E). Additionally, the application of 1μM corticosterone also induced a substantial decrease in the cell surface expression of Ca_V_1.2 channels in cultured hippocampal neurons (Fig. 2F). To further explore the mechanism of glucocorticoids regulating Kv2.1 and Cav1.2 channels, we tested the inhibitory effect of glucocorticoids in heterologous HEK293 cells, which lack endogenous Kv2.1 and Ca_V_1.2 channels(34, 35).

### Glucocorticoids have little effect on Kv2.1 currents but significantly reduce the cell surface expression of Kv2.1 channels in HEK293 cells

The primary glucocorticoids in mice and rats are corticosterone, while in humans, the primary glucocorticoids are cortisol. Therefore, we used cortisol on HEK293 cells transfected with Kv2.1 channels. Extracellular application of 1μM cortisol had little effect on Kv2.1 currents in HEK293 cells (Fig.3A). Similar to the effect of corticosterone on Kv2.1 channels in hippocampal neurons, 1μM cortisol significantly reduced the cell surface expression of Kv2.1 in HEK293 cells (Fig. 3B). Consequently, the level of Kv2.1 channel proteins in the cell cytoplasm increased (Fig. 3C). These results suggest that cortisol caused Kv2.1 endocytosis rather than de-clustering. It is well-known that clustered Kv2.1 channels are nonconducting. Since the effect of cortisol on the surface expression of the Kv2.1 channel is much greater than the effect on the current amplitude, it is reasonable to assume that the endocytosed Kv2.1 channels induced by cortisol are from non-K^+^-conducting Kv2.1 clusters. Therefore, we tested the effect of cortisol on Kv2.1 clusters. We used TIRF microscopy to image the change of Kv2.1 (N-terminal GFP fusion of Kv2.1) clusters in HEK 293 cells. Consistent with previous studies (18), large GFP-Kv2.1 clusters were clearly detected by TIRF (Fig. 3D). Extracellular application of 1μM cortisol quickly reduced Kv2.1 clusters in HEK293 cells (Fig. 3D). Furthermore, we tested the effect of cortisol on mutant Kv2.1S586A channels, which lost the ability to form clusters (36). Extracellular application of 1μM cortisol for 10 minutes did not alter the surface expression of Kv2.1S586A channels (Fig. 3E). The data suggest that glucocorticoids mainly promote the endocytose of Kv2.1 clusters.

**Figure 3.**
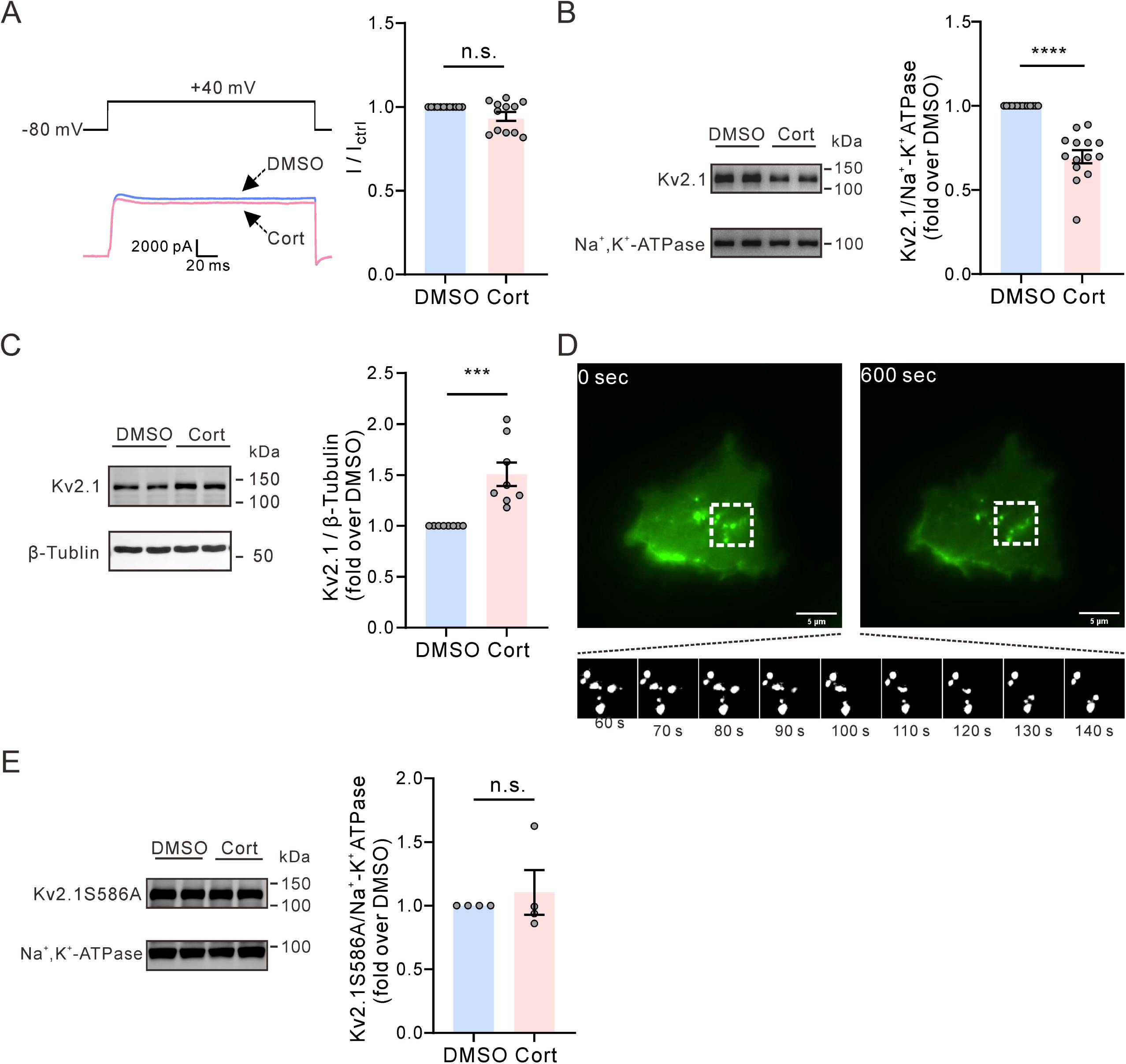
Effect of glucocorticoids on Kv2.1 channels in HEK293 cells. **(A)** Left, representative Kv2.1 current traces induced by a depolarization pulse from −80 to +40 mV under the vehicle control (0.01% DMSO) condition and subsequently in the presence of 1 μM cortisol (Cort) in the same HEK293 cell. Right, statistics for the amplitude of Kv2.1 current from Left using a two-tailed paired *t-*test (n = 12). n.s., not significant (P = 0.059). **(B)** Left, representative western blot of the Kv2.1 channel surface expression after 10-min cortisol treatment (1 μM) in the extracellular solution in HEK293 cells. Right, statistics from 14 independent experiments using a two-tailed unpaired *t-*test. ****P = 2.7E-8. Na^+^-K^+^ ATPase as a membrane protein loading control. **(C)** Left, representative western blot of the cytosol protein level of Kv2.1 channels after 10-min cortisol treatment (1 μM) in the extracellular solution in HEK293 cells. Right, statistics from 8 independent experiments using a two-tailed unpaired *t-*test. ***P = 0.00062. β-tubulin as an internal control. **(D)** Representative TIRF images showing the effect of 1μM cortisol on Kv2.1 clusters in HEK293 cells. **(E)** Left, representative western blot of the Kv2.1S586A mutant channel surface expression after 10-min cortisol treatment in the extracellular solution in HEK293 cells. Right, statistics from 4 independent experiments using a two-tailed unpaired *t-*test. P = 0.5745.

### Glucocorticoids significantly inhibit Ca_V_1.2 currents while has little effect on the surface expression of Ca_V_1.2 channels in HEK293 cells

When compared to its effect on calcium channel currents in hippocampal neurons, glucocorticoids also rapidly reduced the Ca_V_1.2 currents but with smaller amplitude in HEK293 cells (Fig. 4A). Moreover, Cortisol inhibited Ca_V_1.2 currents at all testing potentials that more positive than −10 mV, and did not alter the steady-state activation of Ca_V_1.2 currents (Fig. 4C). Unlike the effect on Ca_V_1.2 in hippocampal neurons, glucocorticoids (1 μM cortisol) did not alter the cell surface expression of Ca_V_1.2 in HEK293 cells (Fig. 4D).

**Figure 4.**
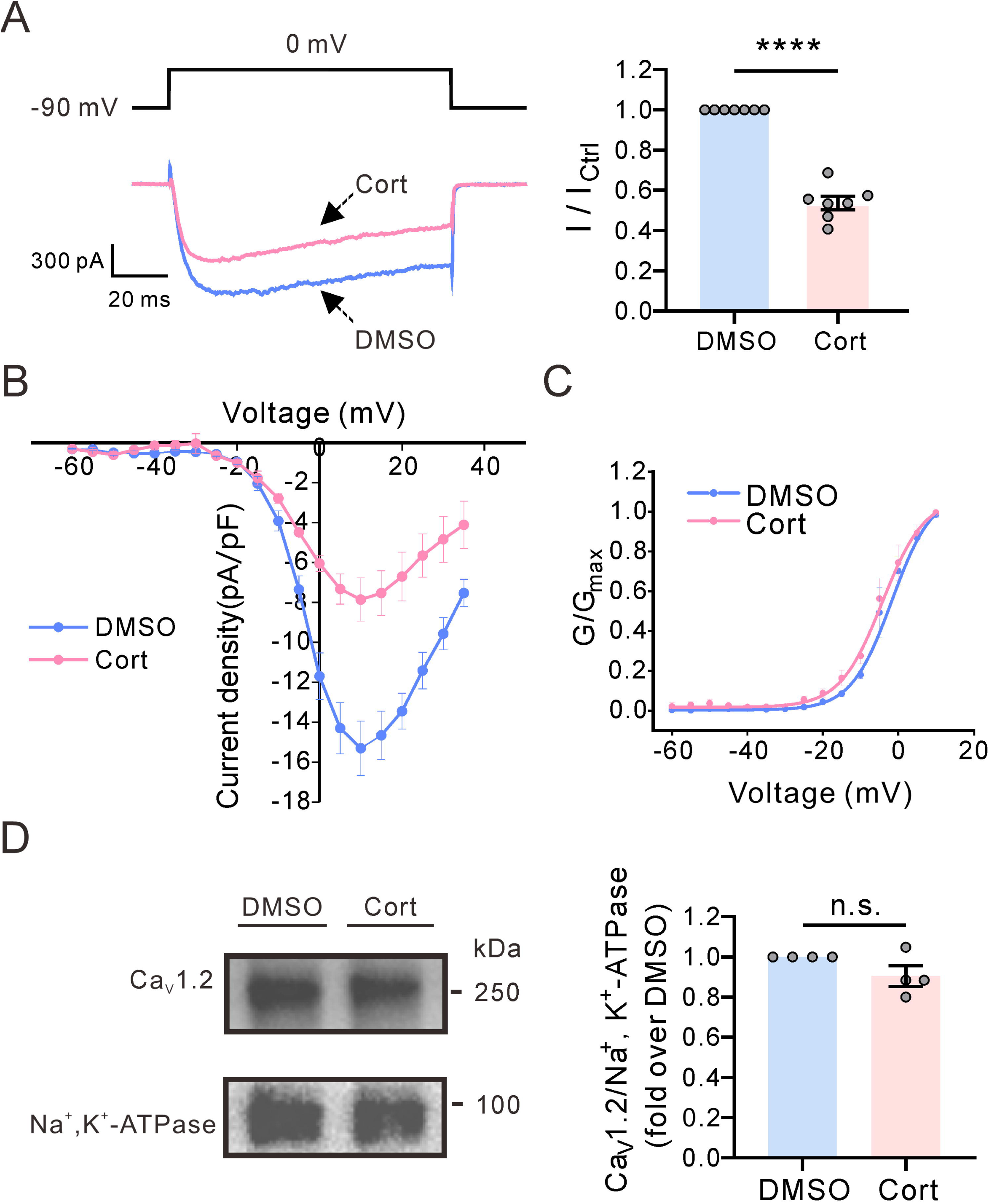
Effect of glucocorticoids on Ca_V_1.2 channels in HEK293 cells. **(A)** Left, representative *I*_Ba_ traces induced by a depolarization pulse from −90 to 0 mV under the control (0.01% DMSO) condition and subsequently in the presence of 1 μM cortisol in the same HEK293 cell. Right, statistics for the amplitude of Kv2.1 current from Left using a two-tailed paired *t-*test (n = 7). ***P = 8.2E-6. **(B)** I-V curve of Ca_V_1.2 currents in the absence or presence of 1 μM cortisol in HEK293 cell. **(C)** 1 μM cortisol did not alter the steady-state activation of Ca_V_1.2 currents. **(D)** Left, representative western blot of the Ca_V_1.2 channel surface expression after 10-min cortisol treatment (1μM) in the extracellular solution in HEK293 cells. Right, statistics from 4 independent experiments using a two-tailed unpaired *t-*test. n.s., not significant (P = 0.1143).

### Glucocorticoids reduce the cell surface expression of Ca_V_1.2 channels when Kv2.1 and Ca_V_1.2 are co-expressed in HEK293 cells

Previous studies have demonstrated a spatial correlation between Kv2.1 clusters and Ca_V_1.2 channels in cultured hippocampal neurons(29, 30). Therefore, we hypothesized that glucocorticoids affect Ca_V_1.2 surface expression through Kv2.1 channel clusters. To test our hypothesis, we co-transfected Ca_V_1.2 with either wild-type Kv2.1 or mutant Kv2.1S586A expression vectors in HEK293 cells. Our results showed that 1 μM cortisol significantly reduced the cell surface express of Ca_V_1.2 channels in the presence of wild-type Kv2.1 channels in the HEK293 cells (Fig. 5A), as seen in cultured hippocampal neurons. Additionally, 1μM cortisol had a greater inhibitory effect on Ca_V_1.2 currents in the co-transfected cells than the cells transfected with Ca_V_1.2 alone (Fig. 5B). Furthermore, we found that 1μM cortisol did not alter the cell surface express of Ca_V_1.2 channels in the presence of mutant Kv2.1S586A channels in HEK293 cells (Fig.5C). This suggests that glucocorticoids induce Ca_V_1.2 channel endocytosis through Kv2.1 clusters.

**Figure 5.**
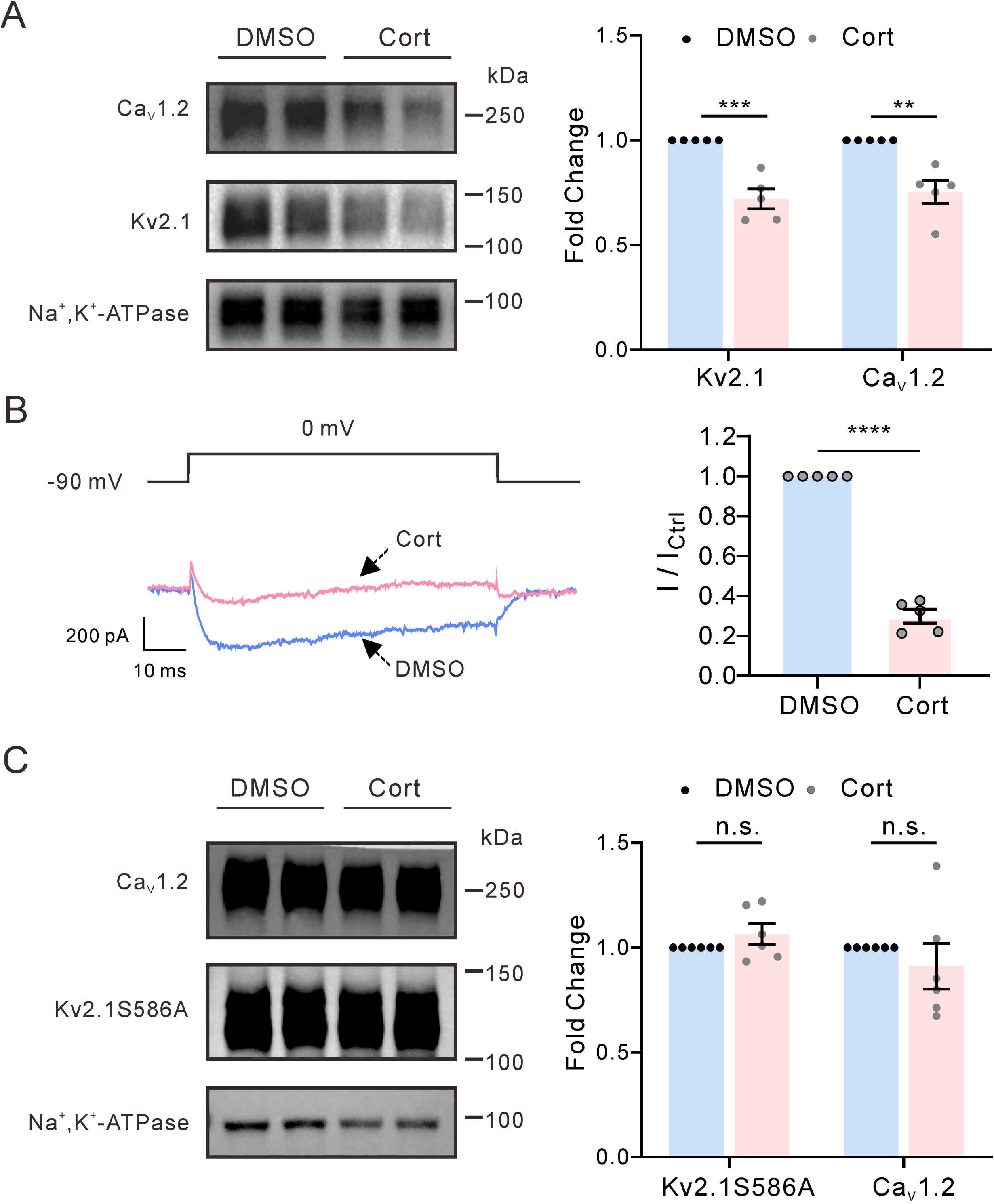
Effect of glucocorticoids on Ca_V_1.2 channels in HEK293 cells expressing Kv2.1. **(A)** Left, representative western blot of the Ca_V_1.2 channel surface expression after 10-min cortisol treatment (1μM) in the extracellular solution in HEK293 cells expressing both Ca_V_1.2 and Kv2.1 channels. Right, statistics from 5 independent experiments using a two-tailed unpaired *t-*test. ***P = 0.0004, **P = 0.00197. **(B)** Left, representative *I*_Ba_ traces induced by a depolarization pulse from −90 to 0 mV under the control condition and subsequently in the presence of 1μM cortisol in the same HEK293 cell. Right, statistics for the amplitude of Kv2.1 current from Left using a two-tailed paired *t-*test (n = 5). ***P = 3.4E-5. **(C)** Representative western blots (left) and statistical analysis (right) show the effect of cortisol on the Ca_V_1.2 channel surface expression when Ca_V_1.2 and Kv2.1S586A channels are co-expressed in HEK293 cells. Two-tailed unpaired *t-*test, n = 6, P = 0.4339.

### Glucocorticoids regulate Kv2.1 channels through membrane-associated receptors

Finally, we investigated how glucocorticoids regulate the membrane expression of Kv2.1 channels. The rapid effect of glucocorticoids is normally mediated by membrane-associated receptors(37). We tested the effect of cortisol-BSA conjugate (cortisol-BSA), which cannot pass through cell membranes, on Kv2.1 channels. We found that 1μM cortisol-BSA had similar effects on the cell surface expression of Kv2.1 as 1μM cortisol in cultured hippocampal neurons (Fig. 6A and B). However, the glucocorticoid receptor inhibitor CORT125281 did not alter the effect of cortisol on Kv2.1 channels (Fig. 6A and B). These data indicate that glucocorticoids regulate cell surface expression of Kv2.1 channels through membrane-associated receptors.

**Figure 6.**
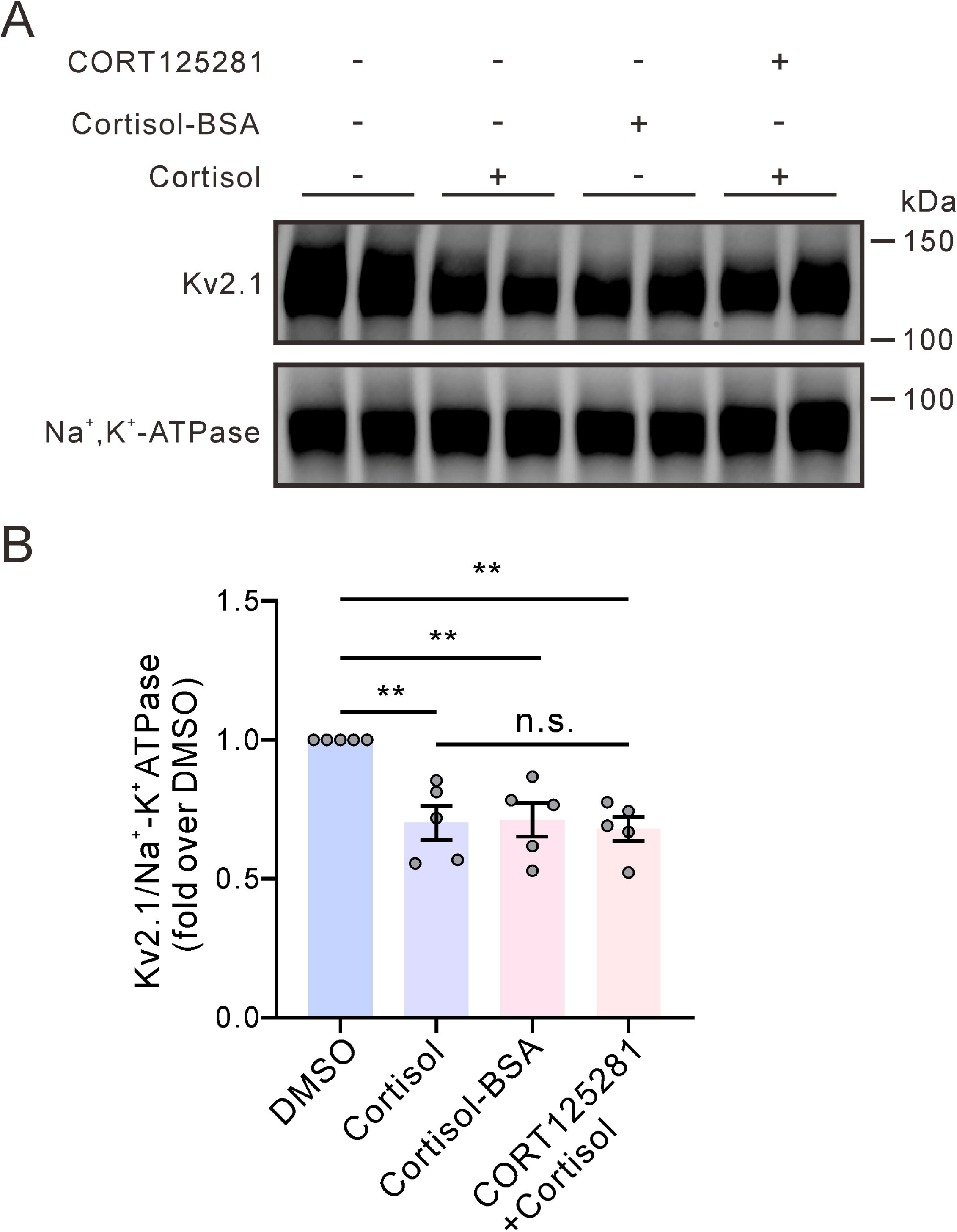
Glucocorticoids modulate the cell surface expression of Kv2.1 channels through membrane-associated receptors. **(A)** Representative western blot images show the effects of cortisol-BSA and cortisol with/without the glucocorticoid receptor inhibitor CORT125281 on Kv2.1 channel surface expression in cultured hippocampal neurons. (**B)** Statistical analysis of the effects of various treatments on the cell surface expression of Kv2.1 using a one-way ANOVA with Bonferroni *post hoc* test (n = 5). **P < 0.01. n.s., not significant.

### Glucocorticoids regulate Kv2.1 and Ca_V_1.2 channels through PKA signaling pathway

Several signaling pathways have been reported for the rapid effects of glucocorticoids through membrane-associated receptors, including PKC, ERK, and especially the cAMP-PKA signaling pathway (37–39). In cultured hippocampal neurons, we observed that 1μM corticosterone rapidly reduced cAMP levels (Fig. 7A) and inhibited PKA activity, as confirmed by a decrease in phosphorylated PKA levels (Fig. 7B). Furthermore, the effect of corticosterone on cell surface expression of Kv2.1 channels in hippocampal neurons was mimicked by PKA inhibitor H89 (1 μM) and abrogated by PKA stimulator forskolin (20 μM) (Fig. 7C).

**Figure 7.**
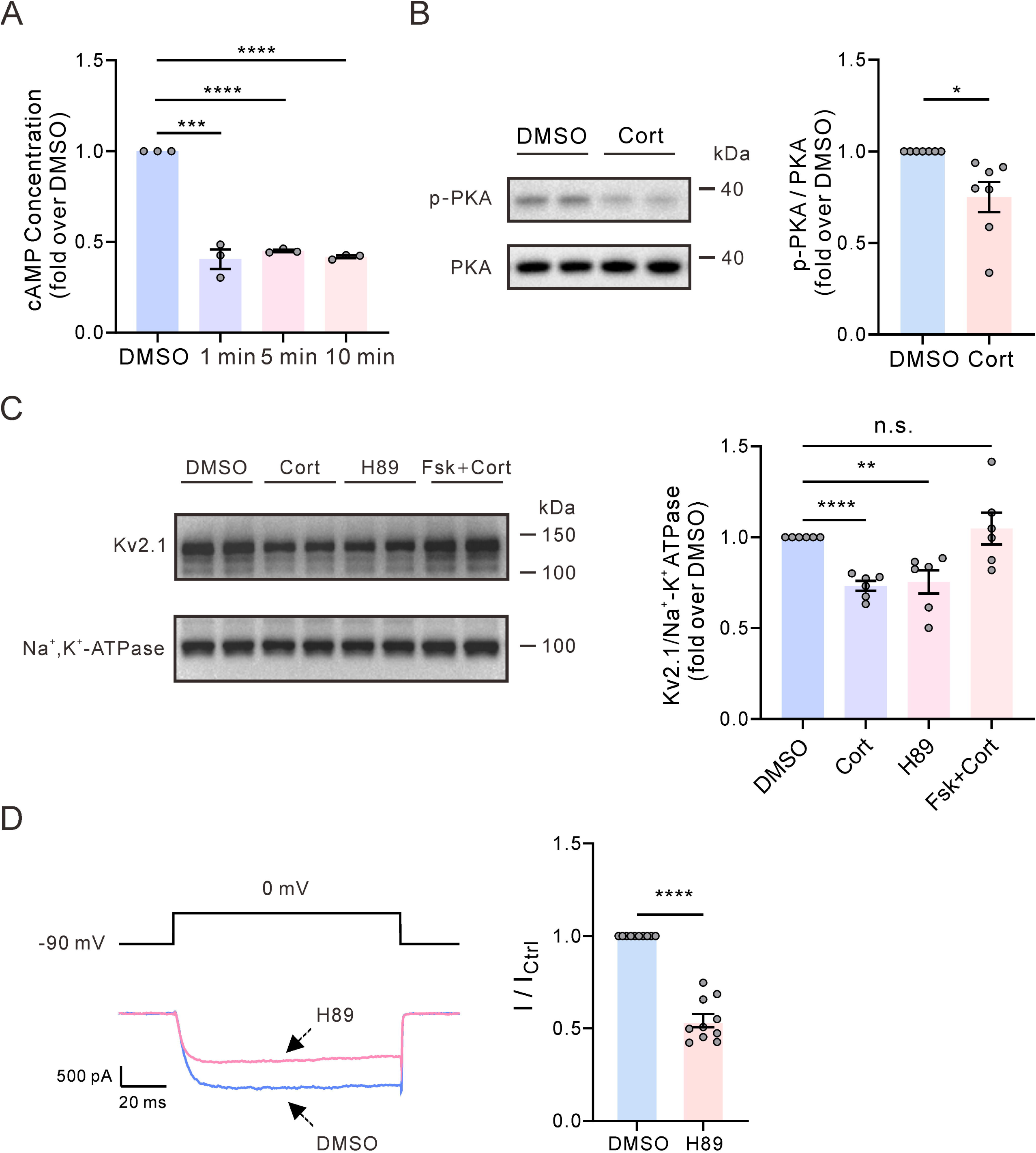
Glucocorticoids reduce the cell surface expression of Kv2.1 and Ca_V_1.2 channels through PKA signaling pathway. **(A)** Extracellular application of 1 μM corticosterone rapidly reduced cAMP levels in cultured hippocampal neurons (n = 3, ***P < 0.001, one-way ANOVA with Bonferroni *post hoc* test). **(B)** Left, representative Western blot of the PKA phosphorylation level after 10-min corticosterone treatment in cultured hippocampal neurons. Right, statistics from 7 independent experiments using a two-tailed unpaired *t*-test. *P = 0.0101. **(C)** Representative western blots (left) and statistical analysis (right) showing the effect of PKA inhibitor H89 and agonist Forskolin (Fsk) on the Kv2.1 channel surface expression in cultured hippocampal neurons. n = 6, ****P < 0.0001, **p = 0.0034, n.s., not significant (p = 0.5948), one-way ANOVA with Bonferroni *post hoc* test. **(D)** Left, representative Ca_V_1.2 current traces induced by a depolarization pulse from −90 to 0 mV under the control condition and subsequently in the presence of 1μM H89 in the same HEK293 cell. Right, statistics for the amplitude of Ca_V_1.2 current from Left using a two-tailed paired *t-*test (n = 10). ****P <0.0001.

Cortisol inhibited Ca_V_1.2 currents but had little effect on the cell surface expression of Ca_V_1.2 in HEK293 cells (Fig. 4), suggesting glucocorticoids regulate Ca_V_1.2 activity besides promoting the channel endocytosis. Previous studies indicate that PKA can directly phosphorylate Ca_V_1.2, and this phosphorylation increases Ca_V_1.2 activity (40). We investigated whether the inhibitory effect of glucocorticoids on Ca_V_1.2 currents was via the PKA signaling pathway. The PKA inhibitor H89 (1 μM) mimicked the effect of cortisol on Ca_V_1.2 currents in HEK293 cells (Fig. 7D).

## Discussion

This study demonstrates that glucocorticoids regulate Ca^2+^ signals in mammalian hippocampal neurons through the PKA signaling pathway in two ways: by inhibiting PKA to reduce Ca_V_1.2 channel activities, and by promoting Kv2.1 clusters endocytosis to increase Ca_V_1.2 channel internalization.

Mammalian brain Kv2.1 channels are typically found in clusters on the somas and proximal dendrites of neurons (41, 42). Increasing evidence suggests that these clustered Kv2.1 channels play important nonconducting roles. Previous studies have shown that Kv2.1 clusters play a structural role in inducing the formation of ER/PM junctions in both cultured hippocampal neurons and transfected HEK293 cells through binding with vesicle-associated membrane protein-associated protein A and B(21, 25). Clustered Kv2.1 channels at ER/PM junctions recruit Ca_V_1.2 to form Ca^2+^ microdomains with ryanodine receptor ER Ca^2+^ release channels, which generates local spontaneous Ca^2+^ release at the soma of hippocampal neurons(29). In this study, we found that glucocorticoids rapidly inhibited spontaneous Ca^2+^ sparks. Glucocorticoids not only inhibit Ca_V_1.2 channel currents, but also disrupt the Ca^2+^ microdomains by targeting clustered Kv2.1 at the soma of cultured hippocampal neurons. The major target of glucocorticoids is clustered Kv2.1 rather than dispersed Kv2.1, as supported by two pieces of evidence. First, glucocorticoids reduced the cell surface expression of Kv2.1 by 30.2%, but had little effect of Kv2.1 currents in HEK293. This can be explained by the fact that clustered Kv2.1 conducts little potassium (43). Second, in the presence of wild-type Kv2.1, glucocorticoids caused a decrease in the surface expression of Ca_V_1.2 channels in HEK293 cells. However, this effect was not observed in the presence of non-clustering Kv2.1S586A mutant channels.

Neuronal Kv2.1 clusters can be de-clustered by stimuli, including kainite, glutamate, hypoxia, and ischemia(32, 44, 45), while none of them affect the total surface expression of Kv2.1. We found that glucocorticoids affect both the clustering status and surface expression of Kv2.1. This suggests a complex underlying molecular mechanism involved. Previous studies showed that the activation properties of Kv2.1 were altered by channel dephosphorylation, but not by actin depolymerization(43). The actin cytoskeleton is important for forming and maintaining Kv2.1 clusters(46, 47), and TTX has been shown to increase Kv2.1 clustering(48, 49) and stabilize the binding between α-actin and Ca_V_1.2(50). We found that preincubation with TTX prevented the glucocorticoid induced endocytosis of Kv2.1 in HEK293 cells (supplemental Fig. S1), suggesting that glucocorticoids may act by changing the cell membrane actin skeleton.

The rapid effects of glucocorticoid are commonly mediated by membrane glucocorticoid receptors. We found cortisol-BSA mimicked the effect of cortisol on Kv2.1, while glucocorticoid receptor antagonist, CORT125281, did alter the effect. This suggests that glucocorticoids inhibit Kv2.1 through unknown membrane receptors. Glucocorticoids rapidly suppress PKA in the AtT20 mouse corticotroph tumor cell line(51), but stimulate PKA in HT4 mouse neuroblastoma cells(52), suggesting a cell-specific effect of glucocorticoids on PKA. PKA phosphorylation significantly influences Ca_V_1.2 activity. Previous studies suggest that AKAP-Anchored PKA is essential for maintaining neuronal Ca_V_1.2 channel activity and NFAT transcriptional signaling (53). Qian et al. reported that β-adrenergic receptor agonist isoproterenol increases Ca_V_1.2 activity by stimulating PKA in hippocampal neurons(54). Here, we found that glucocorticoids decreased cAMP level and PKA activity, thereby inhibiting Ca_V_1.2 currents in hippocampal neurons and HEK293 cells. A recent study showed that glucocorticoids inhibited the levels of cAMP by activation of adhesion G-protein-coupled receptor GPR97(55). Further investigation is needed to determine whether the effect of glucocorticoids reported here is mediated by GPR97.

Glucocorticoids are produced in a diurnal pattern controlled by the hypothalamic-pituitary-adrenal axis. It is now well-known that the glucocorticoid circadian variation is actually composed of an underlying ultradian rhythm. This rhythm occurs less than 60 minutes for rodents(56) and every 60-120 minutes in human(57, 58). Furthermore, the ultradian rhythm of glucocorticoids is not affected by the disruption of circadian inputs to the hypothalamic-pituitary-adrenal activity(59). Recent studies show that the ultradian rhythmicity of glucocorticoids is critical in regulating normal emotional and cognitive responses in humans(4). Wu et al. have revealed that Kv2.1-organized ER/PM junction sites represent around 12% of the somatic surface in mammalian brain neurons(28). At ER/PM junctions, Kv2.1 control somatic Ca^2+^ signals by recruiting Ca_V_1.2, thereby regulating neuronal excitation–transcription coupling(30, 60). Since both Kv2.1 and Ca_V_1.2 are widely expressed in mammalian brain, the rapid effect of glucocorticoids on the Kv2.1-Ca_V_1.2 unit may play an import functional role in the ultradian rhythm of glucocorticoids.

In conclusion, glucocorticoids rapidly regulate Ca^2+^ signals in hippocampal neurons in two ways. First, they inhibit Ca_V_1.2 channel activity (regardless of whether it is coupled with Kv2.1 or not) by reducing PKA activity, which results in a reduction of the overall Ca^2+^ signal. Second, they reduce the surface expression of Ca_V_1.2 channels by promoting the endocytosis of Kv2.1 channels. This disruption of clustered Kv2.1 mediated Ca^2+^ reduces the local spontaneous somatic Ca^2+^ signals that would otherwise regulate excitation-transcription coupling. This study provides further insight into the rapid effect of glucocorticoids in the central nervous system.

## Materials and Methods

### Cell culture and transfection

Human embryonic kidney (HEK293) cells were obtained from the cell bank of the Chinese Academy of Science. These cells were cultured in Dulbecco’s Modified Eagle Medium supplemented with 10% fetal bovine serum and 1% antibiotic antimycotic solution. Plasmids for rat GFP-tagged Kv2.1 channels, and mouse Ca_V_1.2 with β_3_ and α_2_δ_1_ subunits were transiently transfected into HEK293 cells using jetPRIME (Polyplus). For patch clamp recordings, the cells were used after 24 h transfection. Hippocampal neurons from Sprague-Dawley rats of either sex on postnatal day 0 to 1 were used for primary culture. Briefly, hippocampi were dissected and dissociated into single cells with trypsin, and then the cells were plated onto poly-L-lysine–coated glass coverslips inside a six well plate. All experiments were conducted on neurons between 7 and 14 days in vitro (DIV), and were performed on neurons from a minimum of three separate cultures.

All protocols used were approved by the Committee on the Ethics of Animal Experiments of Fudan University, and were in strict accordance with the NIH Guidelines for the Care and Use of Animals.

### Molecular biology

Plasmids for rat GFP-tagged Kv2.1 (34) channels, and mouse Ca_V_1.2(61) were as previously reported. The Kv2.1S586A point mutant was generated using the GFP-Kv2.1 wild-type (WT) plasmid as a template. Site-directed S586A mutagenesis was performed on the Kv2.1 channel using the Quick-Change XL Site-directed Mutagenesis kit (Stratagene, La Jolla, CA, USA). Kv2.1S586A mutation was confirmed by sequencing.

### Live cell imaging

Hippocampal neurons and HEK293 cells (transfected with GFP-tagged Kv2.1) were plated onto poly-L-lysine–coated glass bottom dishes. Imaging was performed in HBSS (14025092, Gibco, NY, USA) solution with 20 mM HEPES (H3375, Sigma-Aldrich, Munich, Germany). Hippocampal neurons were first incubated in regular culture medium with 2 µM fluorescent calcium indicator Cal-590 AM (#20510, AAT Bioquest, CA, USA) for 60 minutes at 37 °C. Dye-containing medium was then aspirated, followed by two washes in HBSS which had been warmed to 37 °C. Cells were then incubated in HBSS for an additional 30 minutes at 37 °C prior to imaging. Images were acquired on a Nikon Eclipse Ti TIRF/widefield microscope equipped with an Andor iXon EMCCD camera and a Nikon LUA4 laser launch with 405, 488, 561, and 647 nm lasers, using a 100x/1.49 NA PlanApo TIRF objective and NIS Elements software. Images were analyzed by Fiji (https://fiji.sc). The Fiji plugin xySpark (62) was used for automated spark detection and analysis.

### Western blot

The surface proteins of HEK293 and hippocampal cells were biotin-labeled and isolated using a Pierce cell surface protein biotinylation and isolation kit (A44390, Thermo Scientific, IL, USA). The protein samples were resolved using 10% SDS PAGE and transferred to polyvinylidene fluoride membranes. The membranes were immunoblotted with anti-Kv2.1 antibody (1:1000, ab192761, Abcam, Cambridge, UK), anti-Ca_V_1.2 antibody (1:1000, ab84814, Abcam), anti-Na^+^-K^+^ ATPase (1:1000, ab76020, Abcam), anti-beta-tubulin antibody (1:2000, ab179511, Abcam), and anti-PKA antibody (1:1000, #4782, Cell Signaling Technology, MA, USA) and anti-phospho-PKA antibody (1:1000, #5661, Cell Signaling Technology). The blots were developed using enhanced chemiluminescence reagents and imaged using either the ChemiDoc XRS+ imaging system from Bio-Rad (Hercules, CA, USA) or the e-BLOT Touch Imager (e-BLOT Life Sciences, Shanghai, China) with manufactureŕs software. Molecular weight marker and chemiluminescent signal images were automatically overlaid by the manufactureŕs software. Quantitative analysis of detected bands was performed using Fiji (https://fiji.sc/).

### Immunofluorescence

Hippocampal neurons on glass coverslips were fixed in 4% paraformaldehyde for 15 minutes at room temperature, washed with cold PBS, and incubated in 0.5% Tween-20 in PBS for 10 minutes. Next, the neurons were blocked with 10% horse serum, 0.3% Triton X-100 in PBS for two hours at room temperature. The primary antibodies used were mouse anti-Kv2.1 (1:500, ab192761, Abcam) and rabbit Ca_V_1.2 (1:500, ab234438, Abcam), and they were incubated with the cells for 1 day at 4°C in a solution containing 1% horse serum and 0.3% PBST. Cells were washed three times with PBS and incubated overnight at 4°C in a secondary antibody solution containing Cy3-labeled Goat Anti-Mouse/Rabbit IgG(H+L) or Alexa Fluor 488-labeled Goat Anti-Mouse IgG(H+L) (1:500, Beyotime, Shanghai, China), 1% horse serum, and 0.3% PBST. Following another wash with PBS, the cells were imaged using the Nikon A1+ Confocal Microscope System (Tokyo, Japan). The images were subjected to rolling ball background subtraction and subsequently converted into a binary mask by thresholding. The sizes of Kv2.1 clusters were measured using the “analyze particles” feature of Fiji.

### cAMP assay

Cultured hippocampal neurons were treated with 1 μM corticosterone for 1, 5 and 10 minutes, respectively. The cells were lysed in 0.1 M HCl. cAMP levels in hippocampal neurons were measured using a cAMP ELISA Kit (NewEast Bioscience, China) following the manufacturer’s instructions.

### Electrophysiology

Whole-cell currents in cultured hippocampal neurons and HEK293 cells were recorded using an Axopatch 200B amplifier (Axon Instruments, Union City, CA, USA) and data were collected with PCLAMP 10.7 software (Axon Instruments). The bath solution for potassium current recording contained (in mM): 140 NaCl, 2.5 KCl, 0.001 TTX and 5 4-AP, 10 glucose, 10 HEPES, and 1 MgCl_2_ (pH adjusted to 7.4 using NaOH). The internal solution contained (in mM): 135 potassium gluconate, 10 KCl, 1 CaCl_2_, 1 MgCl_2_, 10 HEPES, 2 Mg-ATP, and 10 EGTA (pH adjusted to 7.3 using KOH). For calcium current recording, the bath solution contained (in mM): 140 TEACl, 10 BaCl_2_, 0.001 TTX and 5 4-AP, 10 glucose, 10 HEPES, and 2 MgCl_2_ (pH adjusted to 7.4 using TEAOH). The internal solution contained (in mM): 125 Cs-methanesulfonate, 1 MgCl_2_, 10 HEPES, 5 Mg-ATP, 0.3 Tris-GTP and 10 EGTA (pH adjusted to 7.2 using CsOH). The pipettes were made from capillary tubing (BRAND, Wertheim, Germany) and had resistances of 4 to 6 MΩ under these solution conditions. Currents were sampled at 10 kHz and filtered at 2 kHz, and corrected online for leak and residual capacitance transients using a P/4 protocol. All recordings were performed at room temperature.

Cortisol-BSA conjugate (CSB-MC00371b0105) was purchased from CUSABIO (Houston, TX, USA). Cortisol (# B1951), H89 (#B2190), and Forskolin (#B1421) were purchased from APExBIO (Houston, TX, USA). Corticosterone (#HY-B1618) was purchased from MedChemExpress (NJ, USA). All drugs were dissolved in DMSO with a final concentration not exceeding 0.1%.

### Statistical analysis

Data analysis was performed using Clampfit 10.7 (Axon Instruments) and GraphPad Prism (v9.4, GraphPad Software Inc, USA). Normality of the data was checked by the Shapiro-Wilk test. The two-tailed paired or unpaired *t*-test was used to compare two samples, and one-way ANOVA with Bonferroni *post hoc* test was employed for comparing multiple samples. Data are presented as means ± SEM, where ‘n’ indicates the number of tested cells or independent tests. P < 0.05 was considered statistically significant.

## Supporting information

Uncropped western blot images for Figures(incorporated PDF)

Statistical data for all Figures

## Author contributions

Di Wan: Investigation, Formal analysis, Writing – review & editing. Tongchuang Lu: Investigation, Formal analysis, Writing – review & editing. Chenyang Li: Investigation. Changlong Hu: Conceptualization, Supervision, Writing – original draft, Writing –review & editing.

## Acknowledgements

This work was supported d by the National Key Research & Development Program of China (2022YFC3602702), the Science and Technology Innovation 2030 - Brain Science and Brain-Inspired Intelligence Project (2021ZD0201301), the Natural Science Foundation of Shanghai (23ZR1425900), the National Natural Science Foundation of China (31771282).

## Data Availability Statement

All data supporting the findings of this study are available within the paper and its Supplementary Information.

