## Supplemental Figure S1 and supporting text for "Glucocorticoids rapidly modulate Ca_V_1.2-mediated calcium signals through Kv2.1 channel clusters in hippocampal neurons"

***Corresponding author:** Changlong Hu

**This PDF file includes:**

Figure S1
Legends for Movie S1
Legends for Datasets S1 to S2

**Other supporting materials for this manuscript include the following:**

Movie S1
Datasets S1 to S2

A

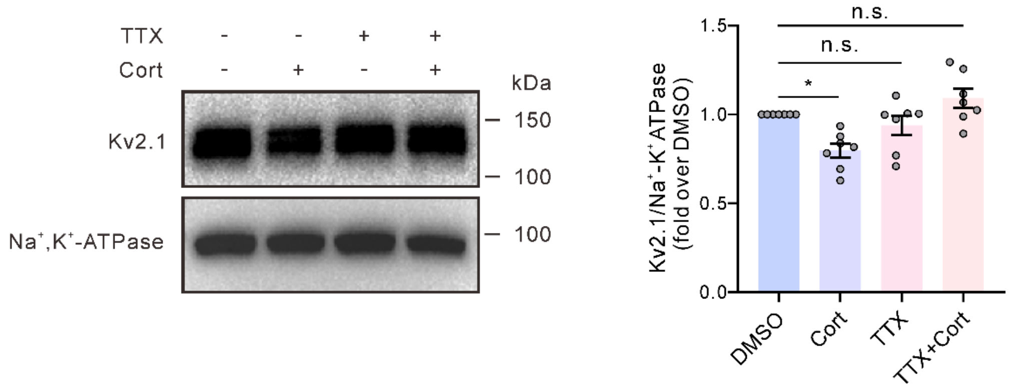


**Figure S1. 1 µM TTX blocked the effect of cortisol on the cell surface expression of Kv2.1 channels in HEK293 cells.**

**(A)** Representative western blots (left) and statistical analysis (right) show the effects of corticosterone and TTX on the Kv2.1 channel surface expression in HEK293 cells. $Na^+$-$K^+$ ATPase as a membrane protein loading control. n = 7. DMSO vs. Cort: *P = 0.0161; DMSO vs. TTX, P > 0.9999; DMSO vs. TTX + Cort: P = 0.9022. n.s., not significant. one-way ANOVA with Bonferroni *post hoc* test.

**Movie S1 (separate file):** A representative video showing the effect of corticosterone (added at around 70 s) on spontaneous somatic $Ca^{2+}$ sparks in hippocampal neurons.

**Dataset S1 (separate file):** The uncropped western blot images for Fig. 2E and F, Fig. 3B and E, Fig. 4D, Fig. 5A and C, Fig. 6A, Fig. 7B and C and Fig. S1.

**Dataset S2(separate file)**: The raw data used for statistical analysis.
