## Supplementary figures and images for "Glucocorticoids rapidly modulate Ca_V_1.2-mediated calcium signals through Kv2.1 channel clusters in hippocampal neurons"

### Uncropped western blot images for Figures(incorporated PDF)

Figure 2E

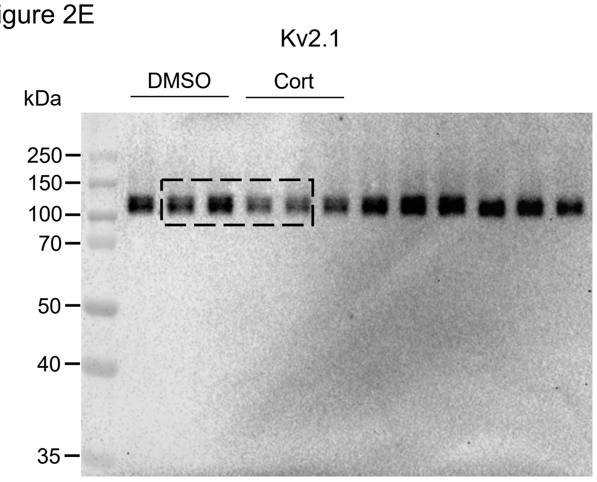


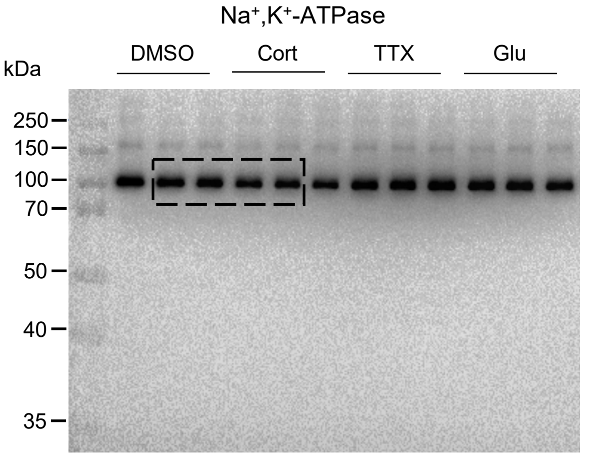


Figure 2F

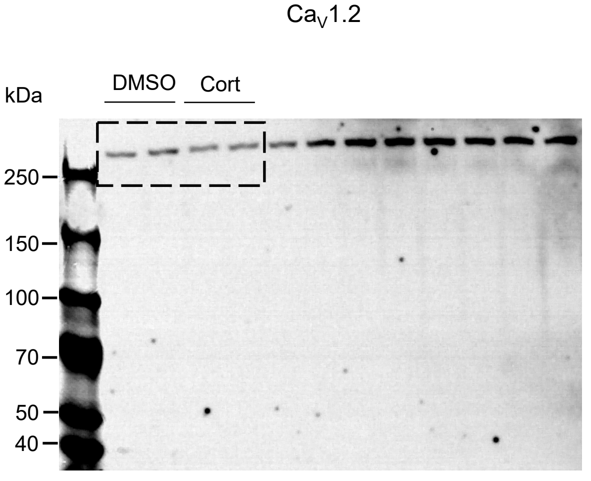


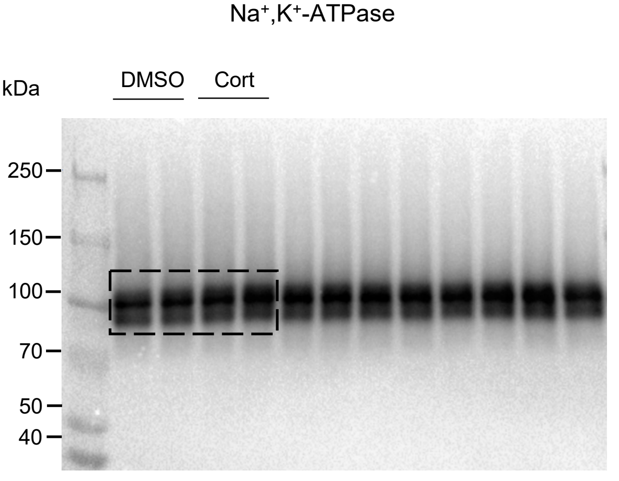

Figure 3B

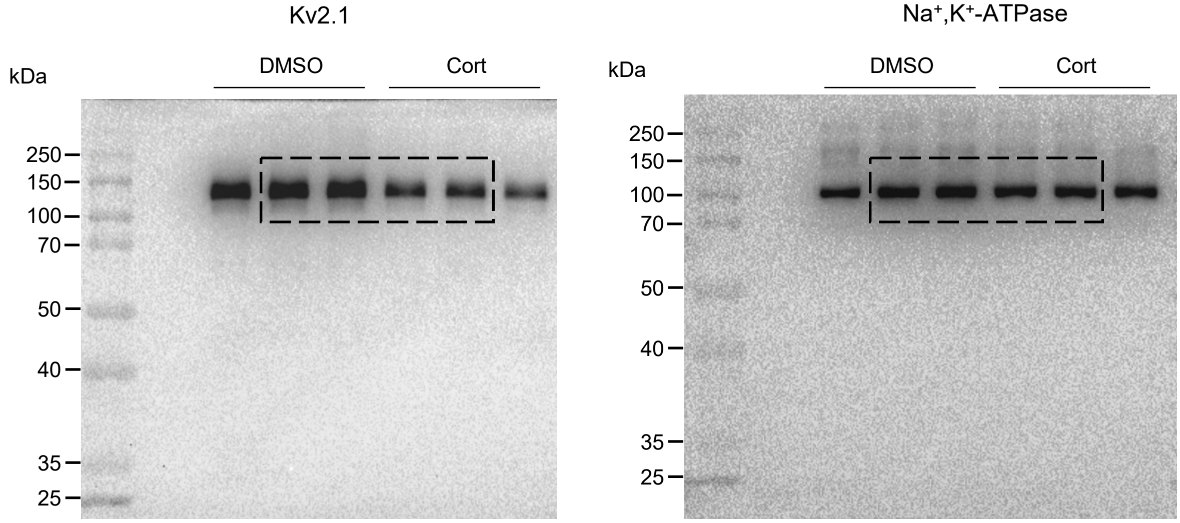


Figure 3C

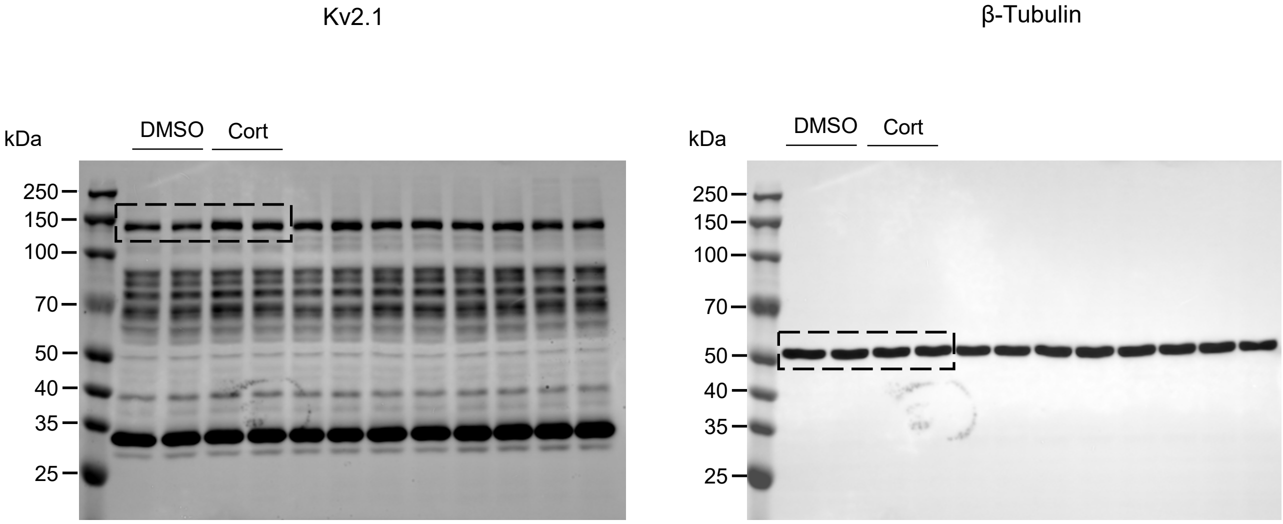


Figure 3E

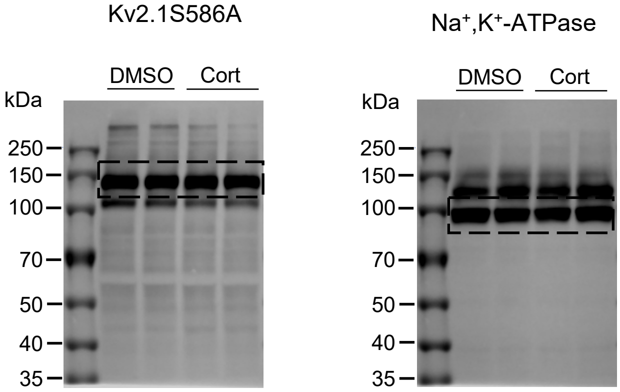

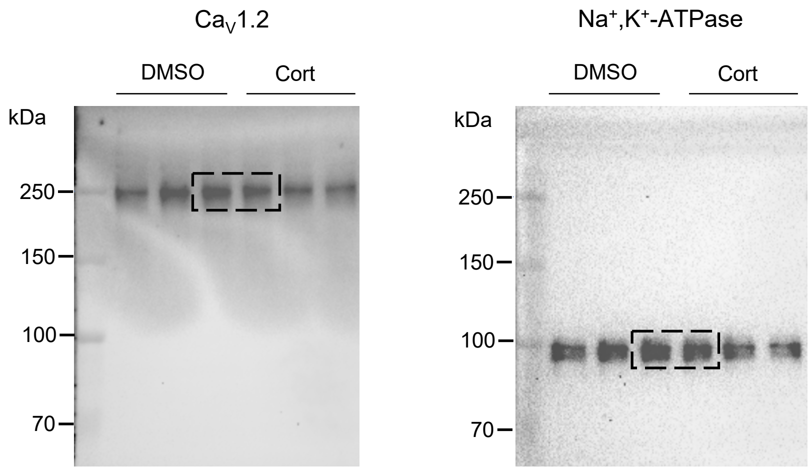

Figure 5A

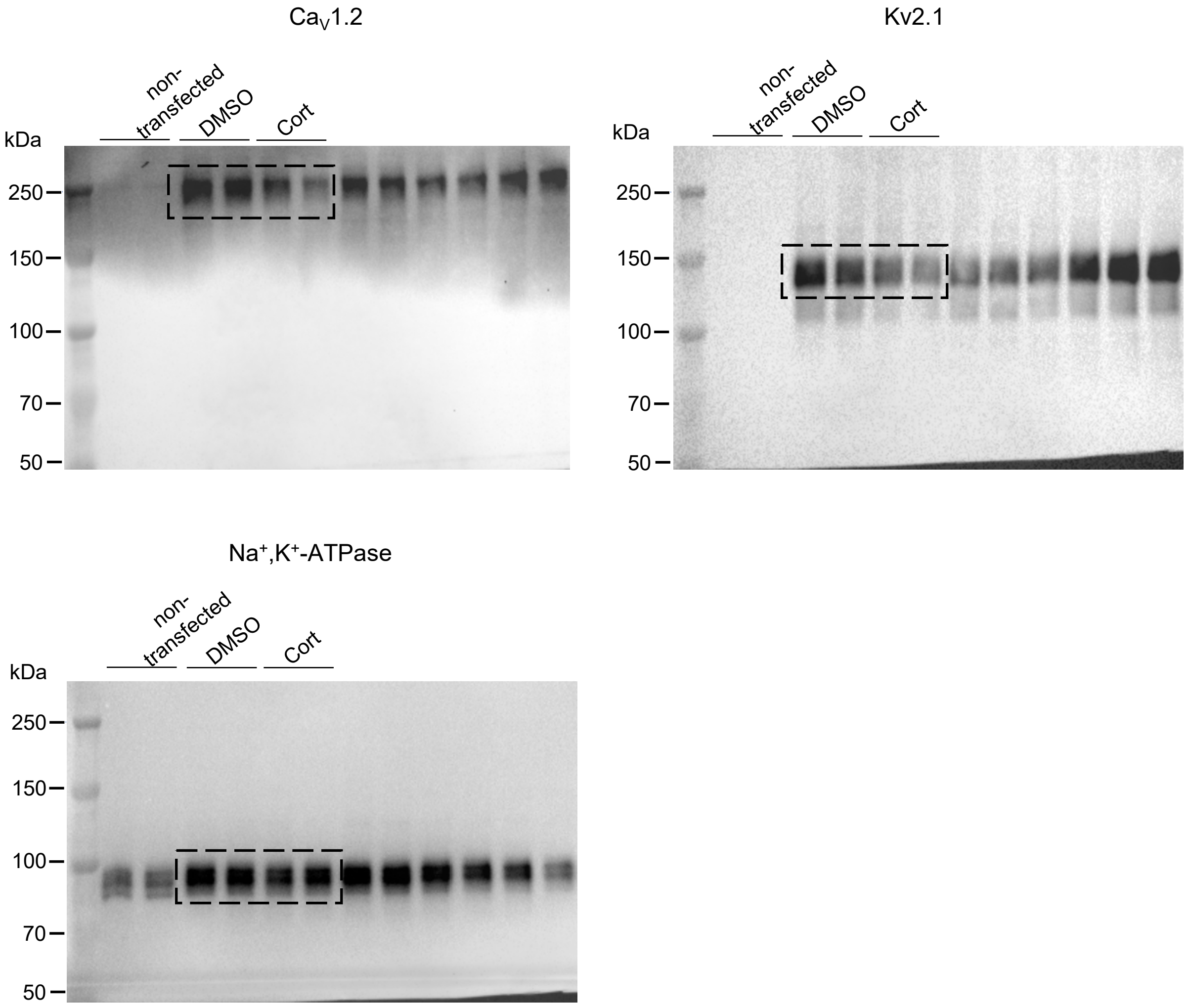


Figure 5C

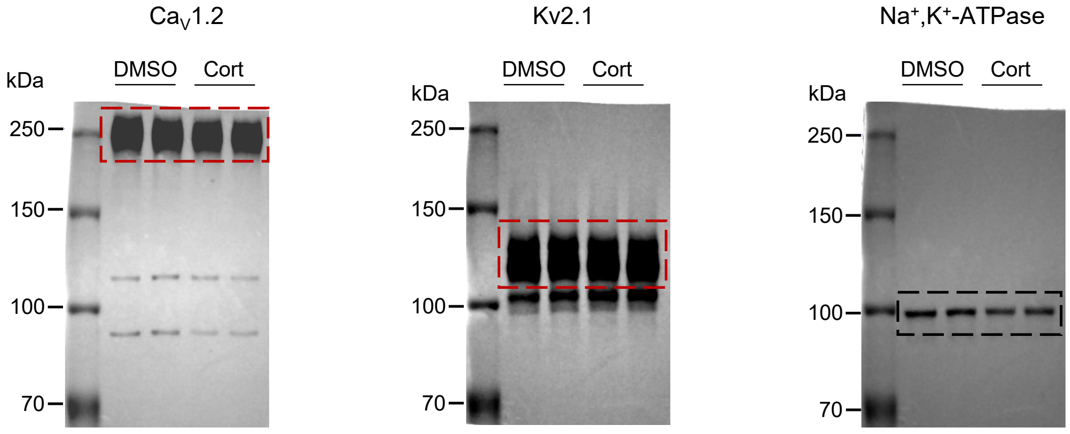

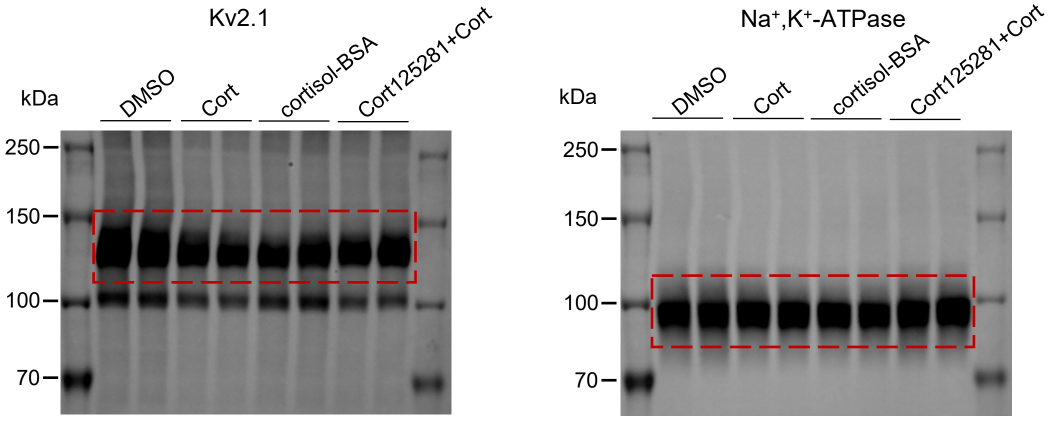

Figure 7B

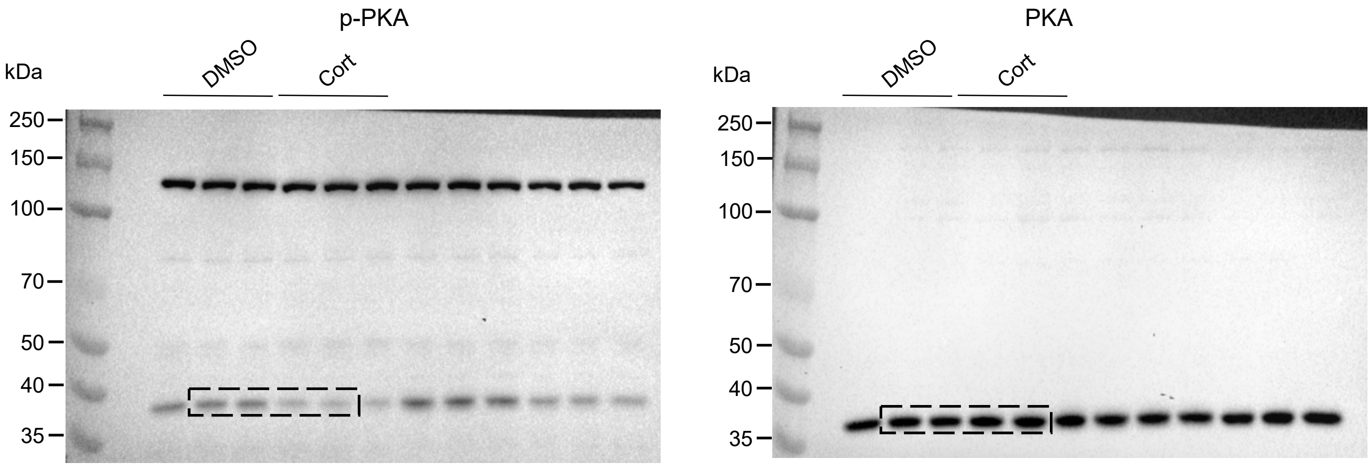


Figure 7C

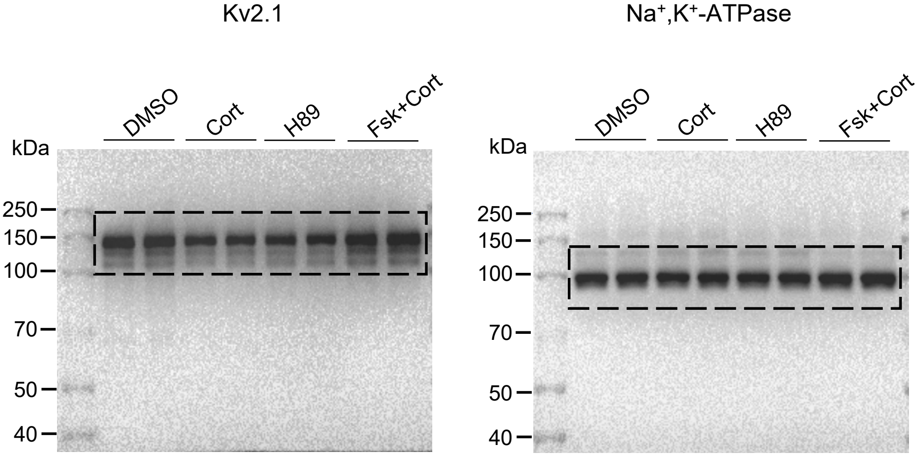

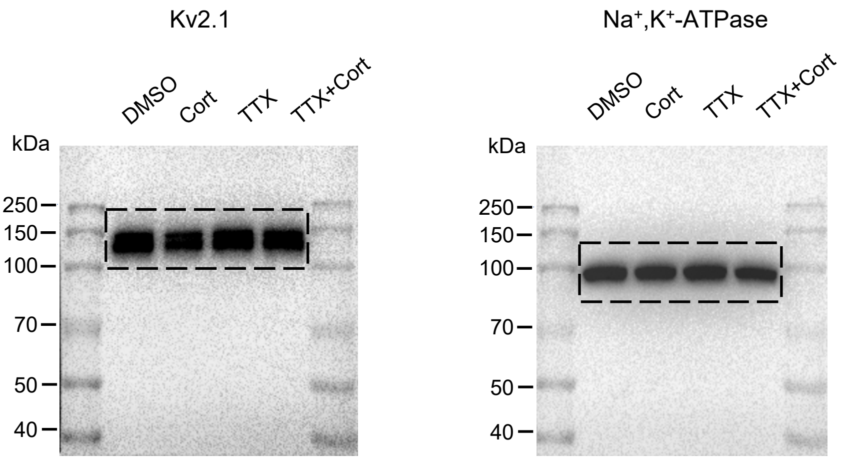
